# *Cdon* is required for organ Left-Right patterning via regulating DFCs migration and the sequential ciliogenesis

**DOI:** 10.1101/2024.03.12.584572

**Authors:** Zhilin Deng, Wenqi Chang, Chengni Li, Botong Li, Shuying Huang, Jingtong Huang, Ke Zhang, Yuanyuan Li, Xingdong Liu, Qin Ran, Zhenhua Guo, Sizhou Huang

## Abstract

*Cdon* and *boc* are members of the cell adhesion molecule subfamily III Ig/fibronectin. Although they were reported to be involved in muscle and neural development at late developmental stage, while their early roles in embryonic development are unknown. Here we discovered that zebrafish *cdon* but not *boc* was expressed in dorsal forerunner cells (DFCs) and epitheliums of Kupffer’s vesicle (KV), implying the possible role of *cdon* in organ LR patterning. Further data showed that the liver and heart LR patterning was disturbed in *cdon* morphants and *cdon* mutants. Mechanically, we found that *cdon* loss of function led to dispersed DFCs migration, smaller KV and defective ciliogenesis, which resulting in randomized *Nodal/spaw* signaling and the sequential organ LR patterning defect. Finally, predominant distribution of a *cdon* MO in DFCs led to defects in DFCs migration, KV morphogenesis/ciliogenesis, *Nodal/spaw* signaling and organ LR asymmetry, being similar to those in *cdon* morphants and *cdon^-/-^* embryos, indicating a cell-autonomous role of *cdon* in regulating KV formation and ciliogenesis during LR patterning. In conclusion, our data demonstrated that, during gastrulation stage and early somitogenesis stage, *cdon* is required for proper DFCs migration, KV formation and ciliogenesis, thus playing an important role in setting up organ LR asymmetry.

## 1. Introduction

Establishment of left-right patterning (LR Patterning) is an important event in early embryonic development, organ asymmetry defect is closely related to the occurrence of various congenital diseases such as congenital heart disease and schizophrenia (Gabriel and Lo, 2020; Huang et al., 2014). Although the mechanisms of establishment of organ LR patterning are complex and diverse in different animal models (Hamada and Tam, 2020; Onuma et al., 2020; Zhu et al., 2020), the role of Kupffer’s vesicle (KV)/Node-Cilia axis in zebrafish and mice is extremely conservative (Forrest et al., 2022; Grimes and Burdine, 2017; Little and Norris, 2021). In zebrafish, the normal development of KV precursor cells (Named dorsal forerunner cells (DFCs) and KV morphogenesis/Ciliogenesis play crucial roles in initiating left-sided signals in the embryonic lateral plate mesoderm (LPM) (Grimes and Burdine, 2017). In organ LR patterning, DFCs proliferate rapidly during gastrulation movement (Abdel-Razek et al., 2023; Gokey et al., 2015), and after bud stage, DFCs differentiate to perform KV and produce cilia in KV, which oscillating to produce a counter-clockwise liquid flow and initiate the specific asymmetric *Nodal/spaw* on the LPM (Bakkers et al., 2009; Liu et al., 2019). Further, the left-sided *Nodal/spaw* and its downstream genes *pitx2* and *lft2* are amplified in the LPM, and the embryonic Midline (such as *shh* and *ntl*) also impedes the expansion of left-sided Nodal signals to the right side of embryo, thus making the left-sided asymmetric Nodal signals more stable (Burdine and Grimes, 2016). Finally at the late stage of organ development, the left-sided Nodal directs a asymmetric migration of organ precursors to establish organ asymmetry patterning (Kajikawa et al., 2022).

During organ LR patterning, the role of zebrafish KV morphogenesis/ciliogenesis and the underlying molecular mechanisms have been received extensive attention. It has been found that most of classical signaling pathways such as Wnt and Fgf (Caron et al., 2012; Neugebauer et al., 2009; Xu et al., 2010; Zhang et al., 2012) and many other genes such as Rab5 (Kuhns et al., 2019) and FOR20 (Xie et al., 2019) participated in KV morphogenesis or ciliogenesis. Our previous studies found that, during zebrafish gastrulation, inhibition of retinoic acid (RA) increased the expression of *fgf8* in DFCs, leading to longer cilia and cilia motility disorders (Huang et al., 2011). More recently, we found that the chemokine receptor *cxcr4a* is highly expressed in DFCs, and the expression of *cxcr4a* promotes CyclinD1 expression to regulate ciliogenesis via regulating the phosphorylation of ERK1/2 (Liu et al., 2019). Although many genes were reported to play crucial roles during KV morphogenesis and ciliogenesis, the mechanisms underlying this process are far from being elucidated.

Cell-adhension molecule-related/down-regulated by oncogenes (Cdon) and its paralog Boc (Brother of Cdo (Kang et al., 2002)) are two members of cell adhesion molecule subfamily III Ig/fibronectin (Jeong et al., 2014; Sanchez-Arrones et al., 2012), they were reported to synergistically or individually regulate different organs development or disease(Jeong et al., 2014; Jeong et al., 2017; Lencer et al., 2023; Sanchez-Arrones et al., 2012; Zhang et al., 2011). As to Cdon, early reports showed that mouse Cdon regulates skeletal muscle and central nervous system development via Shh and N-cadherin/cell adhesion (Cole et al., 2004; Jeong et al., 2014; Zhang et al., 2006a; Zhang et al., 2006b). More detailed studies have shown that Cdon binds to N-cadherin and induces p38/MAPK signals to guide cell differentiation and apoptosis in the process of muscle formation and neural differentiation (Lu and Krauss, 2010). In zebrafish, *cdon* was reported to negatively regulate Wnt signaling pathway during forebrain development to promote neural differentiation and ventral cell fate determination via interacting with LRP6 (Jeong et al., 2014). Further, researchers also found that the localization of Cx43 protein in cells was disorganized in Cdon mutants, resulting in cardiac structural and functional lesions (Jeong et al., 2017). As to Boc, early reports showed that it plays a role in axon guidance and neuron growth (Connor et al., 2005; Okada et al., 2006). Recently, Lencer et al found that in zebrafish *cdon* and *boc* double mutants display trunck neural crest cells (tNCC) migratory defects and loss of slow-twitch muscle, suggesting a non-cell autonomous role of *cdon* and *boc* in regulating tNCC migration (Lencer et al., 2023). Although Cdon and Boc were reported to play important roles in many sections in mouse and zebrafish, their roles in early embryonic development are unknown. During early embryonic development in zebrafish, our data showed that *cdon,* but not *boc,* was expressed in DFCs during gastrulation movement and in epithenial cells of KV at early somitogenesis stage (Fig. 1 and Fig. S1). On the other hand, Cdon has also been reported to control cell apoptosis and cell migration in various kinds of cells (Chapouly et al., 2020; Lencer et al., 2023; Sanchez-Arrones et al., 2012). These facts above suggest that *cdon* may be involved in regulation of clustering movement of DFCs and the sequential organ LR patterning in zebrafish. Here our data identified that *cdon* regulates organ LR pattering via KV/Cilia-*Nodal/spaw* cascade in early embryonic development.

**Figure 1.**
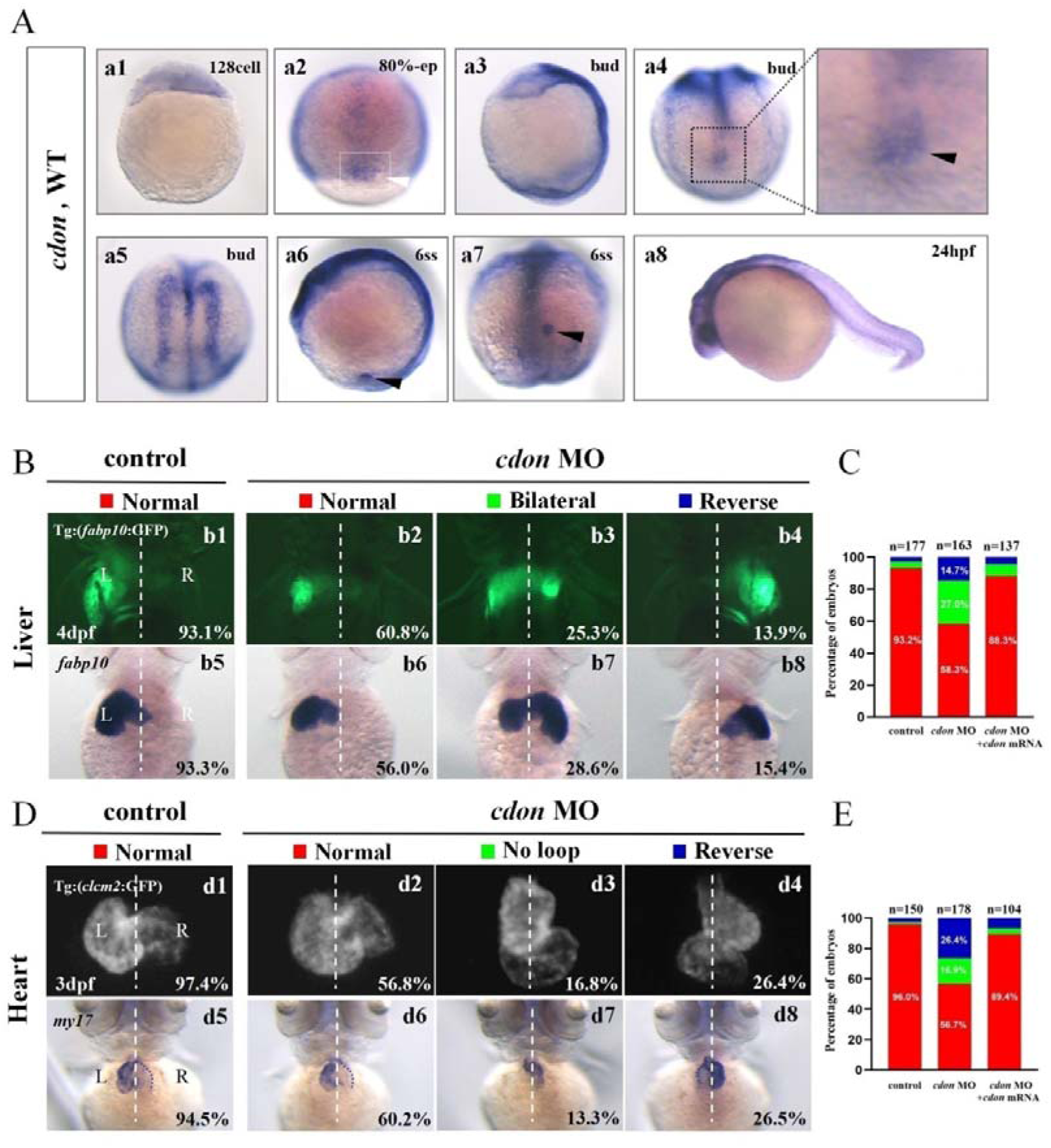
Organ left-right patterning defects in embryos injected with *cdon* MO at 4-cell stage. **(A)** The expression of *cdon* was examined using whole mount *in situ* hybridization from 128-cell stage to 24hpf. *Cdon* was expressed at 128-cell stage (Aa1). White arrow head showed the expression of *cdon* in DFCs at 80% epiboly stage (Aa2). Black arrow head showed the expression of *cdon* in DFCs at bud stage (Aa3-a4). Cdon was also expressed in midline and presumptive neural crest (Aa5). Black arrow head showed the expression of *cdon* in epithelial cell in KV at 6ss (Aa6-a7). Expression of *cdon* was examined at 24hpf (Aa8). **(B)** Embryos injected with *cdon* MO displayed liver LR defects. b1, normal liver in *Tg(fabp10:GFP)* transgenic controls (93.1%, n=102); b2, normal liver in embryos injected with *cdon* MO (60.8%,n=79); b3, liver bifida in embryos injected with *cdon* MO (25.3%,n=79); b4, reversed liver in embryos injected with *cdon* MO (13.9%,n=79). b5, normal liver in wild type controls (93.3%,n=75); b6-b8, embryos injected with *cdon* MO were examined at 4dpf using *in situ* experiments (n=84). **(C)** Percentages of normal liver, liver bifida, and reversed liver in embryos used as control, embryos injected with *cdon* MO and embryos co-injected with *cdon* MO and *cdon* mRNA, in respectively. **(D)** Embryos injected with *cdon* MO displayed heart LR defects. d1, normal-loop in *Tg(cmlc2:GFP)* transgenic controls (97.4%,n=77); d2, normal-loop in embryos injected with *cdon* MO (56.8%, n=95); d3,no loop in embryos injected with *cdon* MO (16.8%, n=95); d4, reversed-loop in embryos injected with *cdon* MO (26.4%, n=95); d5, normal-loop in wild type controls (94.5%,n=73); d6-d8, wild-type embryos injected with *cdon* MO at 4-cell stage were examined for cardiac looping at 72hpf by WISH against *cmlc2* (n=83). **(E)** Percentages of normal looping, no looping, and reversed looping of the heart in embryos being as control, embryos injected with *cdon* MO, and embryos co-injcted with *cdon* MO and *cdon* mRNA. *Cdon* mRNA injection partially rescued the heart LR defect in embryos injected with *cdon* MO (E, the right column showing). Notice: in “C” and “E’, all the transgenic embryos and wild type embryos were used together to calculate the percentage.

## 2 Methods and materials

### 2.1 Zebrafish

Zebrafish wild type (AB), Tg (*cmcl2* :GFP)(Liu et al., 2022), Tg (*fabp10* :GFP)(Liu et al., 2022) and Tg (*sox17* :GFP)(Zhu et al., 2019) lines were maintained at 28.5°C. The embryonic stage was determined according to the external morphology as the standard criteria described before (Kimmel et al., 1995).

### 2.2 Whole-Mount *in situ* hybridization (WISH)

Embryos used for Whole-Mount *in situ* hybridization (WISH) were collected at desired stages and fixed in 4% PFA at 4°C overnight, then the embryos were washed with PBST (2 times, 5min each), dehydrated with MeOH (100%) and stored in MeOH at −20°C.The WISH follows the experimental procedure described previously (Huang et al., 2011; Zhu et al., 2019). The antisense probes *my17*, *fabp10*, *spaw*, *lefty1*, *lefty2*, *sox17*, *sox32* were prepared as previously described (Zhu et al., 2019). To prepare *cdon* antisense probe, the CDs of *cdon* were amplified using PCR, then it was cloned into pCS2+ vector. The detailed process could be find in the section “Plasmid Construction”. Then this construct was used as the template to synthesize *cdon* antisense probe according to previous protocol (Zhu et al., 2019).

### 2.3 Immunostaining

The immunostaining was performed as previous report (Zhu et al., 2019). Briefly, the embryos were fixed with 4% PFA at 4□ overnight, then dehydrated with methanol gradiently and stored at -20□. After methanol /PBST gradient rehydration, PBTN (4% BSA, 0.02%NaN3, PT) was added and the embryos were incubated at 4□ for 3 h, then the α-tubulin antibody (Sigma, T7451, diluted with PBTN at 1:100) was added and incubated overnight on a shaker at 4°C. Next day the embryos were washed with PT (0.3% triton-X-100, in 1X PBS) 4 times (30min each), and the second antibody Goat anti-Mouse IgG(H+L) Alexa Fluor 594 (Sigma, SAB4600105) was added (diluted with PBTN at 1:1000) for overnight incubation (in darkness). Finally, the embryos were rinsed with PT for more than 6 times (30min each) and the imaging was performed.

### 2.4 Plasmids Construction

Total RNA was extracted from zebrafish embryos at 24hpf according to the reagent instructions (TRIzol, Ambion No.15596-026). cDNA was prepared using Revert Aid First Strand cDNA Synthesis Kit (Fermentas No.K1622) according to the manufacturer’s instructions. The CDs of *cdon* was amplified using PCR (Prim STAR Max Premix Takara NO.R045A) and cloned into the vector PCS^2+^ to generate the expression constructs (5x In-Fusion HD Enzyme Premix, Takara

NO.639649). The primers for cloning were as following:

PCS^2+^_F: 5′-CTCGAGCCTCTAGAACTATAGTG-3′,

PCS^2+^_R: 5′-TGGTGTTTTCAAAGCAACGATATCG-3′,

Cdon_F: 5′-TCTTTTTGCAGGATCCGTGAAACAGCGTCATGGAGGAC-3′

Cdon_R: 5′-GTTCTAGAGGCTCGAGTGTTCAGATCTCCTGCACGGT-3′

### 2.5 MO and mRNA injection

The antisense *cdon* morpholino (cdon MOs) was synthesized (Gene Tools) and applied to knock down *cdon* (*cdon* MO (MO^atg^), 5’ -ATAATCTCAGGCCACCGTCCTCCAT-3, 400 uM) (Jeong et al., 2014). *Cdon* MO was injected into the yolk of zebrafish embryos at 1-4 cell stage to block the translation of *cdon* in whole embryos, or at 256-512 cell stage to specifically block the translation of *cdon* mRNA in DFCs. The *cdon* mRNA was synthesized in vitro using mMESSAGE Kit (AM1340, Ambion). In rescue experiments, the concentration for *cdon* mRNA injection was 15ng/ul.

### 2.6 Imaging

Images of the Whole-Mount *in situ* hybridization (in 100% glycerol) were taken with OLYMPUS SZX16 at room temperature. The live embryos of transgenic line Tg (*sox17* :GFP) were placed in 1% Low-Point Melting agrose (LPM agrose) and DFCs were photographed by microscope (OLYMPUS SZX16). To get the images of cilia for immunostaining embryos, the embryos were adjusted with right orientation in 1% LPM agrose and then the cilia were photographed using confocal microscope (OLYMPUS Fluview FV1000).

### 2.7 Statistical analysis

The data were statistically analyzed by Graphpad Prism 9 and ImageJ software. The length of cilia was measured by confocal microscope (OLYMPUS Fluview FV1000). The statistical results were the mean ± SEM of three independent experiments.

### 2.8 Ethics Statement

The study was approved by the Institutional Review Board of Chengdu Medical College (SYXK(□)2015-196), and Zebrafish were maintained in accordance with the Guidelines of Experimental Animal Welfare from Ministry of Science and Technology of People’s Republic of China (2006).

## 3. Results

### 3.1 *cdon* loss of function leads to heart and liver LR patterning defect in *cdon* morphants

In zebrafish, *cdon* and *boc* had been reported to being involved in trunk neural crest cells (NCs) migration and correct proximo-distal patterning in eye development (Jeong et al., 2014; Lencer et al., 2023), demonstrating the crucial role of *cdon* in zebrafish organ development. Since Cdon and Boc synergistically regulate NCs migration (Lencer et al., 2023), facial and eye development (Zhang et al., 2011), to study the role of *cdon* in more early embryonic development, we examined the detailed expression pattern of *cdon* and *boc* from 128-cell stage to 24hpf. The data showed that *cdon* is a maternal factor which distributing in cells unequally (Fig. 1Aa1). During gastrulation movement, beside its expression in the presumptive neural crest and midline (Fig. 1Aa5, (Jeong et al., 2014)), *cdon* was also highly expressed in DFCs (Fig. 1a2, a4). At 6 somite stage, cdon was enriched in the epitheliums of KV (Fig. 1A5-6). On the contrary, even though *boc* was maternally and ubiquitously expressed during gastrulation (Fig. S1A, B), but we did not find *boc* was enriched in DFCs and epitheliums of KV (Fig. S1D-F). Since disturbing DCFs development and the sequential KV/Ciliogenesis leads to organ LR patterning defects (Xie et al., 2019; Zhu et al., 2019), we proposed the possibility that *cdon* is involved in organ LR patterning. To preliminarily and rapidly evaluate this hypothesis, an ATG MO for *cdon* was used to down-regulate the function of *cdon* (Jeong et al., 2014), then organ LR patterning was examined. The results showed that, in *Tg(fabp10:GFP)* transgenic embryos, injection of *cdon* MO did not lead to embryonic defect (Fig. S2), but gave rise to liver LR patterning defect (Fig. 1Bb1-b4, C). Part of embryos injected with *cdon* MO displayed liver bifida (Fig. 1Bb3) and right-sided liver (Fig. 1Bb4). Being similar, in wild type embryos, injection of *cdon* MO also led to liver LR patterning defect (Fig. 1Bb5-b8, C). Next, we examined whether injection of *cdon* MO lead to heart looping defect. The results showed that injection of *cdon* MO caused heart LR patterning defect (Fig. 1D, E), many of *cdon* morphants displayed linear heart (Fig. 1Dd3,d7, E) and reversed heart looping (Fig. 1D4,8, E). To further confirm the role of *cdon* in liver and heart LR patterning, finally we evaluated whether injection of *cdon* mRNA could rescue the organ LR patterning defect in *cdon* morphants. The data showed that injection of *cdon* mRNA partially restored organ LR patterning defect in embryos injected with *cdon* MO (Fig. S3 and Fig. 1C, E; the right columns showed). These data above indicated that *cdon* plays a critical role during organ LR patterning in zebrafish.

### 3.2 left-sided *Nodal/spaw* cascade is randomized in *cdon* morphants

The crucial role of *Nodal/spaw* in LR patterning was proved in previous studies (Kang et al., 2013; Raya and Izpisua Belmonte, 2006; Speder et al., 2007), disruption of *Nodal/spaw* leads to organ laterality defect (Huang et al., 2011). While in some kinds of mutants with organ LR patterning defect, *Nodal/spaw* was not disturbed (Huang et al., 2014; Kurpios et al., 2008; Trinh and Stainier, 2004; Yin et al., 2010). To reveal how *cdon* regulates organ laterality, we examined whether the left-sided *Nodal/spaw* was disturbed in embryos injected with *cdon* MO. The data showed that the left-sided *Nodal/spaw* was disturbed in *cdon* morphants, displaying both-sided, right-sided and disappeared *spaw* expression (Fig.2Aa2-a4, B). Furthermore, we sequentially checked *lefty1* and *lefty2*, the downstream genes of *spaw* in *cdon* morphants. The data showed that the left-sided expression pattern of *lefty2* and *lefty1* in the heart field were also perturbed (Fig. 2Cc2-c4,D, Ee2-e4.F). Since *cdon* was also expressed in midline (Fig. S1Aa5, (Jeong et al., 2014)), we observe whether lefty1 in midline was affected. The data showed that the expression of *lefty1* in middle line was not affected (Fig. 2 Cc2-c4, red arrow showed). These data above suggested that the left-sided *Nodal/spaw* cascade was disturbed in *cdon* morphants, which may lead to organ LR patterning defect in *cdon* morphants.

**Figure 2.**
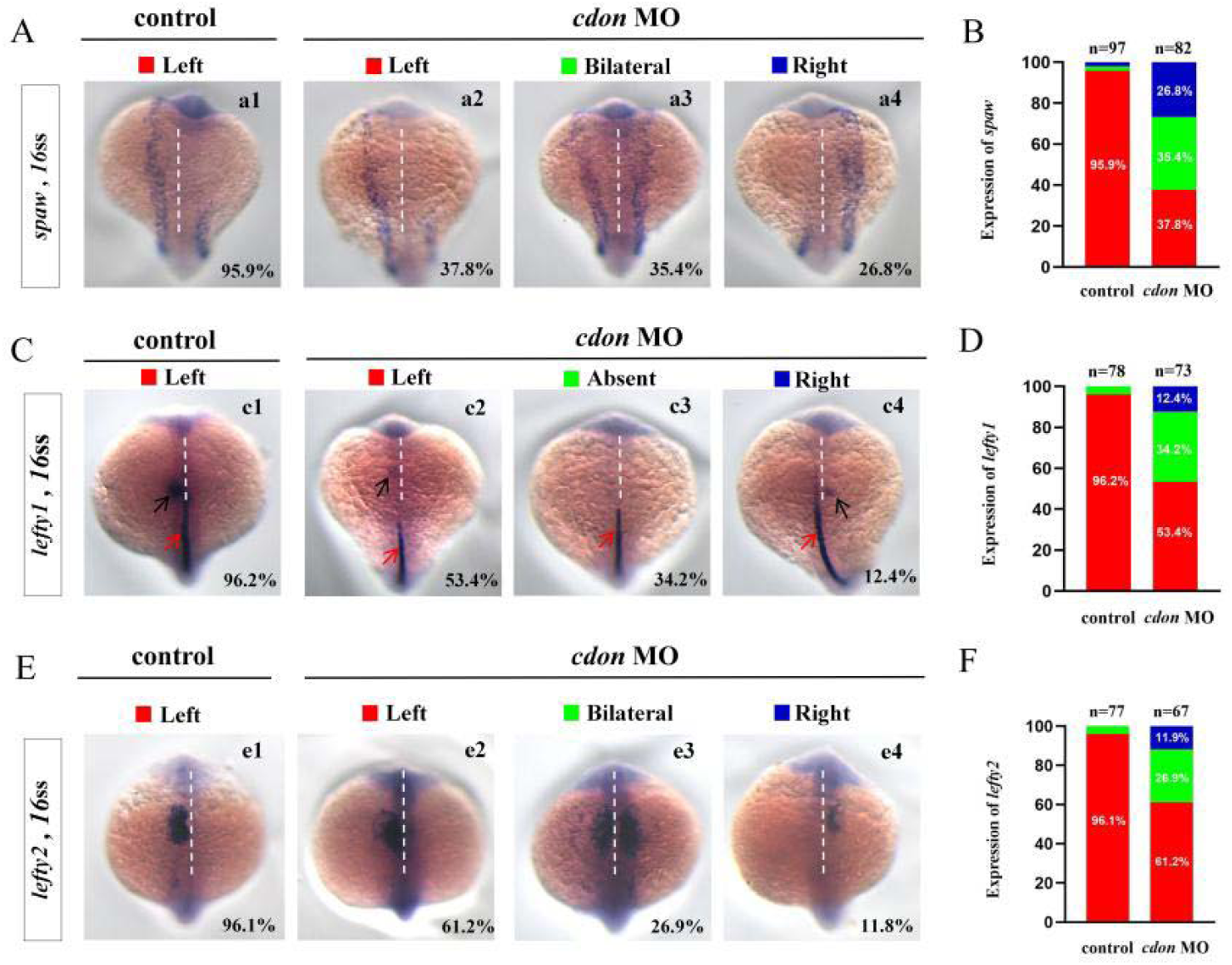
Expression of left-sided Nodal signaling in controls and embryos injected with *cdon* MO. **(A)** Expression of *spaw* was evaluated in different group of embryos. a1, left-sided *spaw* in controls (95.9%, n=97); a2, left-sided *spaw* in embryos injected with *cdon* MO (37.8%, n=82); a3, bilateral *spaw* in embryos injected with *cdon* MO (35.4%, n=82); a4, right-sided *spaw* in embryos injected with *cdon* MO (26.8%, n=82).(**B)** Statistical analysis was performed for the expression of *spaw* in controls and embryos injected with *cdon* MO. **(C)** *Lefty1* is expressed in the heart field (black arrow) and trunk midline (red arrow) in controls and embryos injected with *cdon* MO. c1, left-sided *lefty1* in embryos being as controls (96.2%,n=78, black arrow showed); c2, left-sided *lefty1* in embryos injected with *cdon* MO(53.4%, n=73, black arrow showed); c3, absent *lefty1* in embryos injected with *cdon* MO (34.2%,n=73); c4, right-sided *lefty1* in embryos injected with cdon MO (12.4%,n=73). **(D)** Statistical analysis was performed for the expression of *lefty1* in controls and embryos injected with *cdon* MO. **(E)** *Lefty2* is examined in control embryos and embryos injected with *cdon* MO. e1, left-sided *lefty2* in controls (96.1%, n=77); e2, left-sided *lefty2* in embryos injected with *cdon* MO (61.2%, n=67); e3, bilateral expression of *lefty2* in embryos injected with *cdon* MO (26.9%,n=67); e4, right-sided *lefty2* in embryos injected with *cdon* MO (11.8%,n=67). **(D)** Statistical analysis was performed for the expression of *lefty2* in controls and embryos injected with *cdon* MO.

### 3.3 Clustering DFCs migration, KV morphogenesis and ciliogenesis are disturbed in *cdon* morphants

Our current research showed that *cdon* was expressed in the DFCs at gastrulation stage and in the epithelial cells of KV (Fig. 1Aa2-a7). In zebrafish, DFCs will form the KV at early somite stage, as well the defective KV morphogenesis or defective ciliogenesis will give rise to disturbed left-sided *Ndoal*/*spaw* and the subsequent organ LR defect (Essner et al., 2005; Long et al., 2003; Wang et al., 2011). To find out whether *cdon* regulates DFCs development, KV morphogenesis or ciliogenesis, we analyzed the expression of DFCs marker *sox17* and *sox32*, KV morphogenesis and ciliogenesis in *cdon* morphants. Compared with that in control embryos, at 80% epiboly stage, the clustering DFCs migration was disturbed in *cdon* morphants (Fig. 3 Aa2-a3, Aa5-a6, B). At 10-13 SS the KV was smaller in majority of *cdon* morphants (Fig.3 Cc3, D), and a small part of morphants displayed tiny/absent KV (Fig. 3Cc4, D). Further, we examined the cilia development and found that *cdon* morphants displayed mild shorter cilia than that in controls (Fig.3 Ee2-e4, F), the cilia number was also decreased in *cdon* morphants (Fig.3 Ee2-e4, G). These results showed that the normal expression of *cdon* is required for DFCs migration, KV morphogenesis and ciliogenesis from gastrulation stage to early somitogenesis stage.

**Figure 3.**
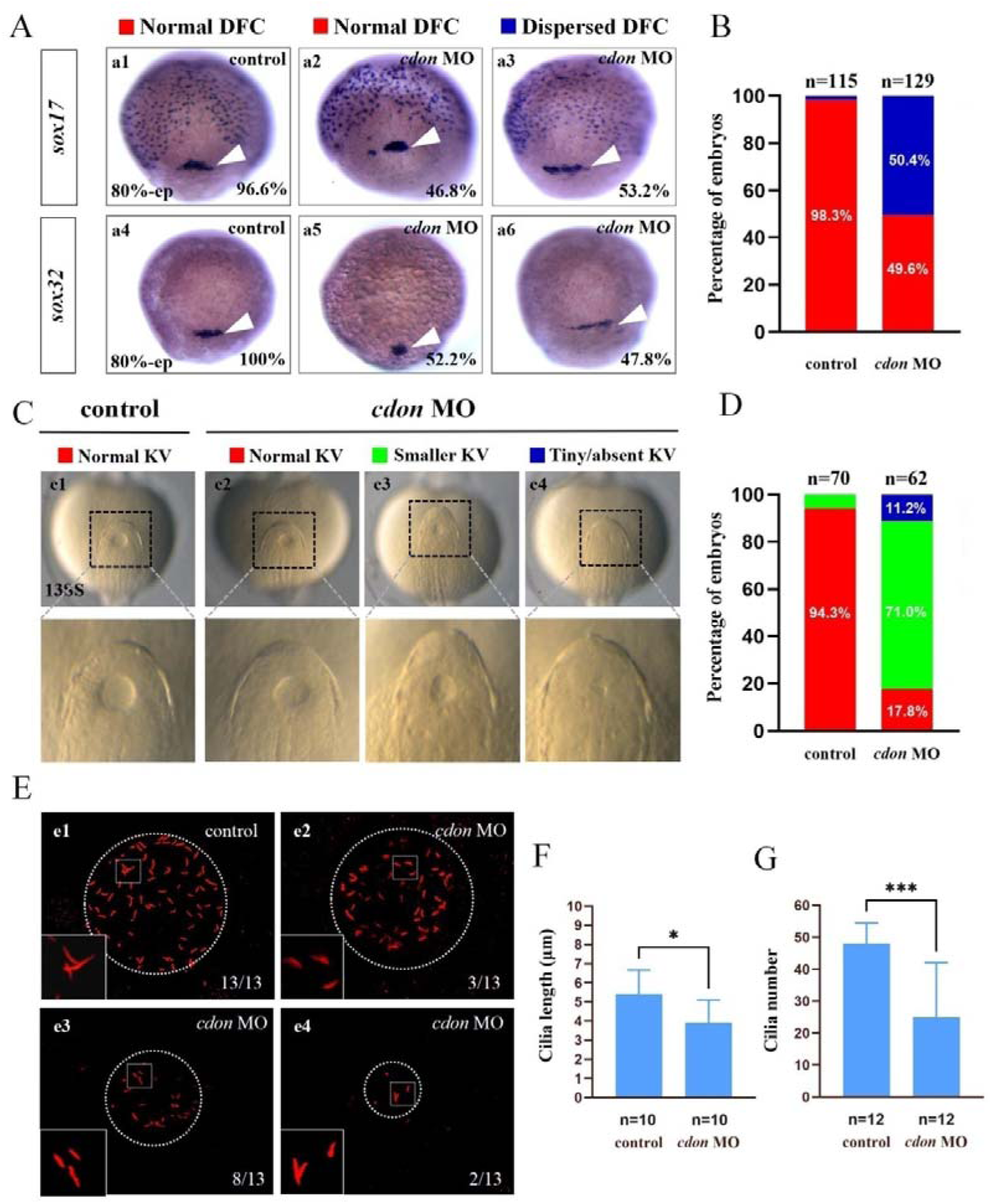
*Cdon* loss of function disturbs clustering DFCs migration, KV morphogenesis and ciliogenesis. **(A)** Expression of *sox17* and *sox32* was examined using WISH at 80% epiboly (white arrow). a1, normal expression of *sox17* in controls (96.6%, n=58); a2, normal expression of *sox17* in embryos injected with *cdon* MO at 4-cell stage (46.8%, n=62); a3, dispersed expression of *sox17* in embryos injected with *cdon* MO at 4-cell stage (53.2%, n=62). a4, normal expression of *sox32* in controls (100%, n=57); a5, normal expression of *sox32* in embryos injected with *cdon* MO at 4-cell stage (52.2%, n=67); a6, dispersed expression of *sox32* in embryos injected with *cdon* MO at 4-cell stage (47.8%, n=67). **(B)** Statistical analysis was performed for the expression of *sox17* and *sox32* in controls and embryos injected with *cdon* MO. Here all the embryos staining with *sox17* or *sox32* were used together to calculate the percentage. **(C, D)** KV morphology was evaluated. c1, normal KV in embryos in controls (94.3%, n=70); c2, normal KV in embryos injected with *cdon* MO at 4-cell stage (17.8%, n=62); c3, smaller KV in embryos injected with *cdon* MO at 4-cell stage (71.0%, n=62); c4, tiny/absent KV in embryos injected with *cdon* MO at 4-cell stage (11.2%, n=62). **(E)** Number and length of cilia were evaluated in controls and embryos injected with *cdon* MO. e1, cilia in embryos being as controls; e2-e4, cilia in embryos injected with *cdon* MO at 4-cell stage. **(F)** Statistical chart for cilia length in KV. **(G)** Statistical chart for cilia number in KV. “*”p<0.05,“***”p < 0.001.

### 3.4 *Cdon* mutation leads to organ LR patterning defect

To confirm the role of *cdon* in organ LR patterning, we generated a *cdon* mutant line using CRISPR-Cas9 method (Shankaran et al., 2017). To generate the *cdon* mutant, we selected a specific sequence in the exon3 of *cdon* as the target sequence (Fig. 4A). As the result, in F1 adults we screened out a frame shift mutation line (Fig. 4B, C). In this mutation, the sequence "AAGGGC" in the exon3 of *cdon* gene was changed to "TTGATGAATGGGGG" (Fig. 4B, C), which resulting in a truncated Cdon protein (only 104 amino acids) (Fig. 4C). In addition, even though we found the expression of *cdon* mRNA was greatly downregulated in *cdon^-/-^*embryos at 8 somite stage and 24hpf (Fig. S4C, D), the *cdon^-/-^* embryos have no distinct external phenotype at different stages (Fig. S4 A, B) and could grow up to adults. Then we evaluated whether this frame shift mutation leads to liver and heart LR patterning defect using *in situ* experiments. The data showed that *cdon^-/-^* embryos, but not control embryos (Fig. 4Dd1), displayed liver LR patterning defect: 71.2% of *cdon^-/-^*embryos displayed left-sided liver (Fig. 4Dd2, E), 17.8% of *cdon^-/-^*embryos displayed bilateral liver (Fig. 4Dd3, E), 11.0% of *cdon^-/-^*embryos displayed right-sided liver (Fig. 4Dd4, E). Being similar, there is no heart LR patterning defect was observed in control embryos (97.5%; Fig. 4Ff1, G), but *cdon^-/-^* embryos displayed no-loop heart (16.4%; Fig. 4Ff3, G) and reversed loop heart (10.9%; Fig. 4Ff4, G). To further confirm the critical role of *cdon* in organ LR patterning, we examined whether injection of *cdon* mRNA could rescue liver and heart LR patterning defect in *cdon^-/-^* embryos. The data showed that injection of *cdon* mRNA partially restored liver and heart LR patterning (Fig. 4E, G). All these data in *cdon^-/-^*embryos further demonstrated that *cdon* is essential for organ LR patterning.

**Figure 4.**
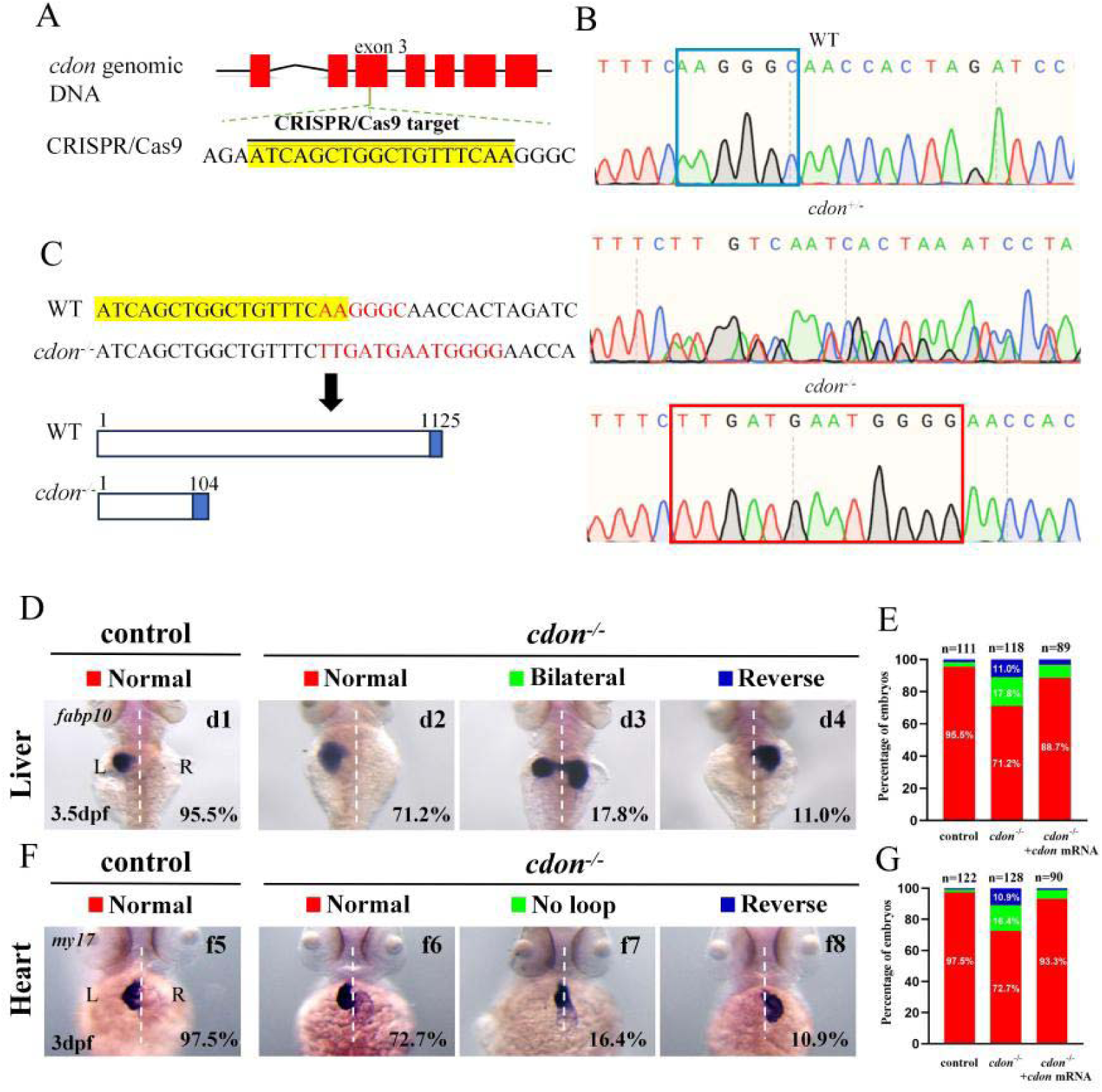
Organ LR patterning defects in *cdon^-/-^*embryos. **(A)** CRISPR/Cas9 target was showed. The sequence in the exon3 of *cdon* gene was chose as the target (yellow highlighted). (**B)** Sequencing results for the genomic DNA in WT embryos, *cdon^+/-^* embryos and *cdon^-/-^* embryos. **(C)** Nucleotide sequences highlighted in yellow are CRISPR/Cas9 target. In the *cdon* mutant, the sequence "AAGGGC" was changed to "TTGATGAATGGGGG". *Cdon* mutant harbors a frameshift mutation that is predicted to result in the production of truncated Cdon protein (104 amino acids). **(D)** *Cdon^-/-^* embryos were found to display liver LR defects using WISH. d1, left-sided liver in controls (95.5%, n=111); d2, left-sided liver in *cdon^-/-^* embryos (71.2%, n=118); d3, liver bifida in *cdon^-/-^*embryos (17.8%, n=118); d4, right-sided liver in *cdon^-/-^* embryos (11.0%, n=118). **(E)** Percentages of left-sided liver, liver bifida, and right-sided liver in *cdon^-/-^*embryos (n=118), controls (n=111) and *cdon^-/-^* embryos injected with *cdon* mRNA (n=89). **(F)** *Cdon^-/-^* embryos displayed heart LR defects. f1, normal-loop heart in controls (97.5%, n=122); f2, normal-loop heart in *cdon^-/-^* embryos (72.7%, n=128); f3, no loop heart in *cdon^-/-^*embryos (16.4%, n=128); f4, reversed loop heart in *cdon^-/-^* embryos (10.9%, n=128). **(G)** Percentages of normal looping, no looping, and reversed looping of the heart in*cdon^-/-^* embryos (n=128), controls (n=122) and *cdon^-/-^*embryos injected with *cdon* mRNA (n=90).

### 3.5 KV/cilia-*Nodal/spaw* cascade was also disturbed in *cdon* mutants

To further confirm the mechanism how *cdon* regulates organ LR patterning, we also examined whether KV/cilia-*Nodal/spaw* cascade was affected in *cdon^-/-^* embryos. First, we examined the expression of *sox17* and *sox32* at 80% epiboly stage to evaluate whether the clustering DFCs migration is disturbed in *cdon^-/-^* embryos. The data showed that 55.7% of *cdon^-/-^*embryos displayed dispersed expression of *sox17* (Fig. 5Aa3, B), 48.0% of *cdon^-/-^*embryos displayed dispersed expression of s*ox32* (Fig. 5Aa6, B). On the contrary, the expression of *sox17* and *sox32* was normal in control embryos (Fig. 5Aa, Aa4, B). This data indicated that DFCs migration was disturbed in *cdon^-/-^*embryos. Then we evaluated whether KV morphogenesis and ciliogenesis were affected in *cdon^-/-^*embryos. The data showed that, in many *cdon^-/-^*embryos, the KV became smaller or absent (Fig. 5 Cc3-c4, D). Being similar to that in *cdon* morphants, in *cdon^-/-^*embryos the cilia length is mild shorter (Fig. 5Ee1-e4, F) and cilia number is decreased (Fig. 5Ee1-e4, G). Finally, we examined the expression pattern of *spaw* in controls and *cdon^-/-^* embryos. The data showed that the expression of left-sided *spaw* was also disturbed in *cdon^-/-^* embryos, displaying left-sided *spaw* (Fig. 5Hh2, I), bilateral *spaw* (Fig. 5Hh3, I) and right-sided *spaw* (Fig. 5Hh4, I). In addition, the expression of *Nodal/spaw* downstream gene *lefty1* and *lefty2* was also disturbed in *cdon^-/-^*embryos (Fig.S5A-D). These data above further confirmed that KV/cilia-*Nodal/spaw* cascade may mediate *cdon* to regulate heart and liver LR patterning during early development.

**Figure 5.**
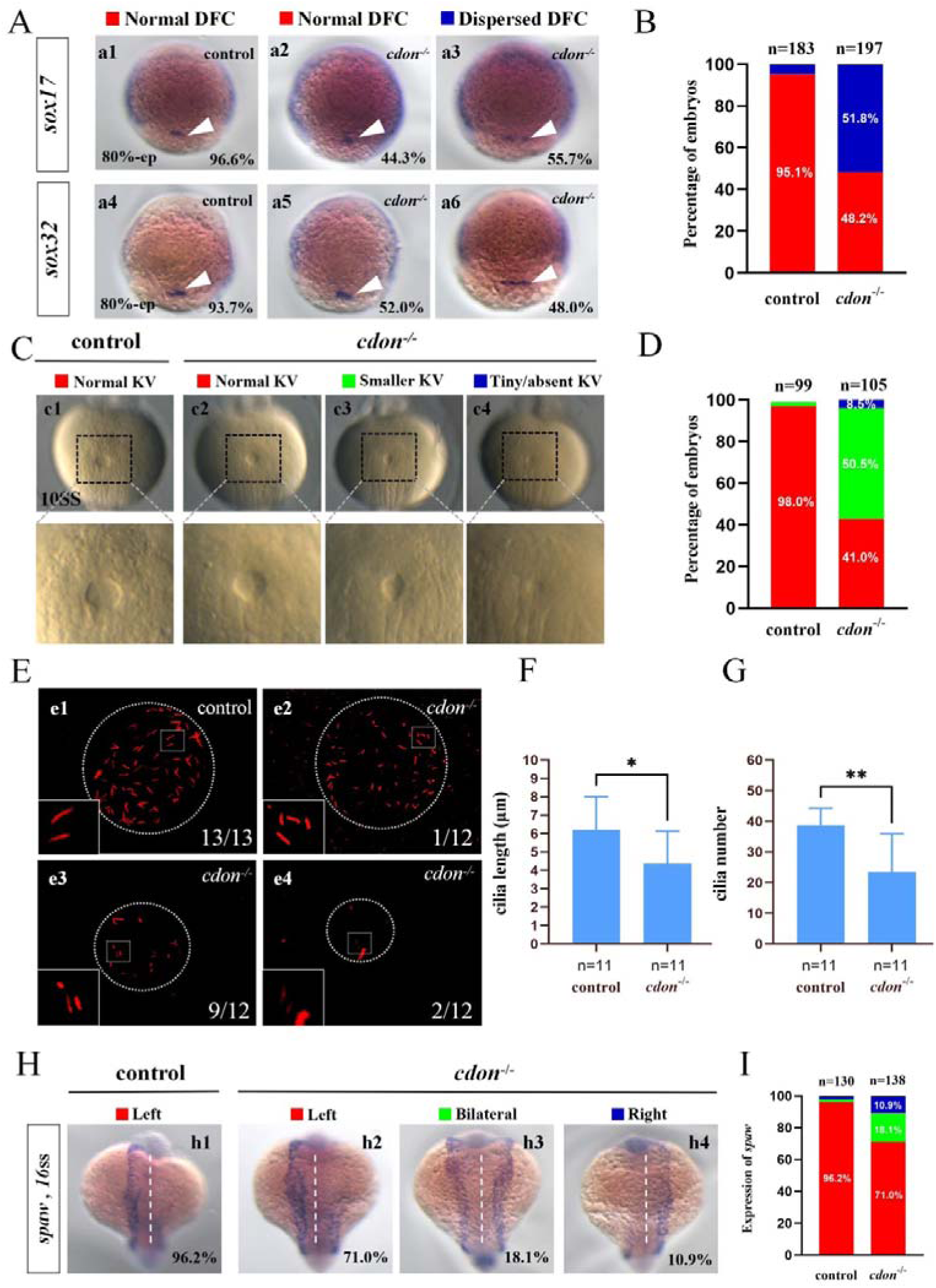
*Cdon* mutation gave rise to defects in KV formation and ciliogenesis. **(A)** Expression of *sox17* and *sox32* was examined using WISH at 80% epiboly. a1, normal expression of *sox17* in controls (96.6%, n = 88); a2, normal expression of *sox17* in *cdon^-/-^*embryos (44.3%, n = 97); a3, dispersed expression of *sox17* in *cdon^-/-^*embryos (55.7%, n = 97). DFCs (white arrow). a4, normal expression of *sox32* in controls (93.7%, n = 95); a5, normal expression of *sox32* in *cdon^-/-^*embryos (52.0%, n = 100); a6, dispersed expression of *sox32* in *cdon^-/-^*embryos(48%, n =100). DFCs (white arrow). **(B)** Statistical analysis was performed for DFCs clustering migration in controls (n=183) and *cdon^-/-^*embryos(n=197). Here all the embryos staining with *sox17* or *sox32* were used together to calculate the percentage. **(C)** Morphology of KV in different group of embryos at 10ss. c1,normal KV in controls (98.0%, n =99); c2,normal KV in *cdon^-/-^*embryos (41.0%, n =105); c3,smaller KV in *cdon^-/-^*embryos (50.5%, n = 105); c4, tiny/absent KV in *cdon^-/-^*embryos (8.5%, n =105). **(D)** Statistical analysis was performed for KV morphogenesis in controls (n=99) and *cdon^-/-^*embryos (n=105). **(E)** Thenumber and the length of cilia were examined at 10ss. e1, cilia in controls; e2-e4, cilia in *cdon^-/-^*embryos. **(F)** Statistical chart for cilia length in KV. **(G)** Statistical chart for cilia number in KV. “*”p<0.05,“**”p < 0.01. **(H)** Expression of *spaw* in controls and *cdon^-/-^* embryos. h1, left-sided *spaw* in controls (96.2%, n=130); h2, left-sided *spaw* in *cdon^-/-^* embryos (71.0%,n=138); h3, bilateral *spaw* in *cdon^-/-^* embryos (18.1%, n=138); h4,right-sided *spaw* in *cdon^-/-^*embryos (10.9%, n=138). (I) Statistical analysis was performed for the expression of *spaw* in controls (n=130) and *cdon^-/-^* embryos (n=138).

### 3.6 *Cdon* loss of function in DFCs results in organ LR patterning defects

Our data have showed that *cdon* was not only expressed in DFCs and KV epithelial cells, but also expressed in some other cells such as cells in midline and PSM (Fig. 1A and Fig. S4 C, D). To confirm whether *cdon* regulates organ LR patterning via DFCs-KV/cilia-*Nodal/spaw* cascade specifically, we injected *cdon* MO at 256-512 cell stage to predominantly block translation of *cdon* mRNA in DFCs (Amack and Yost, 2004; Zhu et al., 2019), then evaluated whether organ LR patterning was disturbed. Indeed, in the embryos injected with *cdon* MO at 256-512 cell stage, the liver and heart LR patterning was disturbed (Fig. 6A-D): In both *Tg(fabp10:GFP)* transgenic embryos (Fig. 6Aa1-a4, B) and wild type embryos (Fig. 6Aa5-a8, B), after predominantly down-regulating the function of *cdon* in DFCs, the livers were localized in left side, both side and right side, in respectively. In both *Tg(cmlc2:GFP)* transgenic embryos (Fig. 6Cc1-c4, D) and wild type embryos (Fig. 6Cc5-c8, D), after predominantly down-regulating the function of *cdon* in DFCs, the hearts displayed normal loop, linear and reversed loop, in respectively. Next, we evaluated whether DFCs-KV/cilia cascade was affected after predominantly down-regulating the function of *cdon* in DFCs. The data showed that the clustering DFCs migration was disturbed (Fig. S6Aa1-a6, B), the size of KV was smaller (Fig. S6 Cc1-c4, D), the length of cilia was mild shorter (Fig. S6Ee1-e4, F) and the number of cilia was also decreased (Fig. S6Ee1-e4, G). Finally, we continued to evaluate if *Nodal/spaw* signaling was disturbed in embryos injected with *cdon* MO at 256-512 cell stage. As result, the expression of left-sided *Nodal/spaw* and its downstream gene *lefty1* and *lefty2* was also randomized in embryos injected with *cdon* MO at 256-512 cell stage (Fig. 6E-J): To the expression of *spaw*, 44.6% (Fig. 6 Ee2), 40.2% (Fig. 6Ee3) and 15.2%(Fig. 6Ee4) of embryos injected with *cdon* MO displayed left-sided *spaw*, both-sided *spaw* and right-sided *spaw*, in respectively; while 97.5% of control embryos displayed left-sided *spaw* (Fig. 6Ee1). To the expression of *lefty1*, 54.9% (Fig. 6Gg2), 29.7% (Fig. 6Gg3) and 15.4% (Fig. 6Gg4) of embryos injected with *cdon* MO displayed left-sided *lefty1*, disappeared *lefty1* and right-sided *lefty1*, in respectively; while 96.5% of control embryos displayed left-sided *lefty1*. To the expression of *lefty2*, 60.9.6% (Fig. 6Ii2), 27.6% (Fig. 6Ii3) and 11.5% (Fig. 6Ii4) of embryos injected with *cdon* MO displayed left-sided *lefty2*, both-sided *lefty2* and right-sided *lefty2*, in respectively, while 97.2% of control embryos displayed left-sided *lefty2* (Fig. 6Ii1). These data indicated that the left-sided *Nodal/spaw* signaling was randomized after down-regulating the function of *cdon* in DFCs. In conclusion, all these data above suggested that *cdon* specifically regulates organ LR patterning via DFCs-KV/cilia-*Nodal/spaw* cascade.

**Figure 6.**
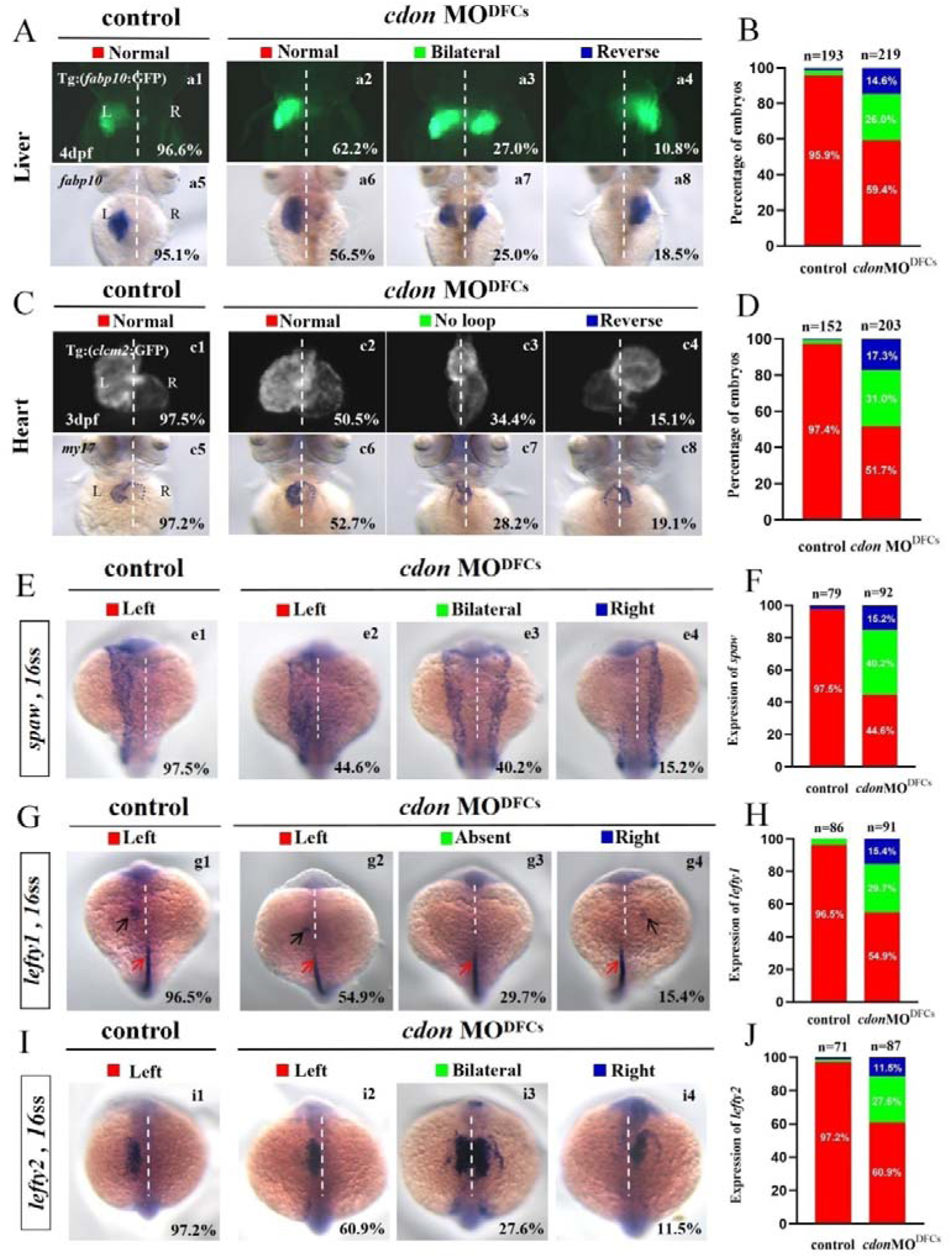
Organ left-right patterning defects in embryos injected with *cdon* MO at the 256-cell stage. **(A)** Embryos injected with *cdon* MO at 256-cell stage was found to cause liver LR defects. a1, normal liver in *Tg(fabp10:GFP)* transgenic controls (96.6%, n=90); a2, normal liver in *Tg(fabp10:GFP)* transgenic embryos injected with *cdon* MO at the 256-cell stage (62.2%, n=111); a3, liver bifida in *Tg(fabp10:GFP)* transgenic embryos injected with *cdon* MO at the 256-cell stage (27.0%, n=111); a4, reversed liver in *Tg(fabp10:GFP)* transgenic embryos injected with *cdon* MO at the 256-cell stage (10.8%,n=111). a5, normal liver in wild type controls (95.1%, n=108); a6-a8,wild-type embryos injected with *cdon* MO at the 256-cell stage were examined for liver laterality at 4d by WISH against *fabp10* probe (n=108). **(B)** Percentages of normal liver, liver bifida, and reversed liver in control embryos and embryos injected with *cdon* MO at the 256-cell stage.**(C)** Heart morphogenesis in *Tg(cmcl2:GFP)* transgenic embryos and wild-type embryos injected with *cdon* MO at the 256-cell stage. c1, normal-loop in *Tg(cmcl2:GFP)* transgenic controls (97.5%, n=80); c2, normal-loop in *Tg(cmcl2:GFP)* transgenic embryos injected with *cdon* MO at 256-cell stage (50.5%, n=93); c3, no loop in *Tg(cmcl2:GFP)* transgenic embryos injected with *cdon* MO at 256-cell stage (34.4%, n=93); c4, reversed-loop in *Tg(cmcl2:GFP)* transgenic embryos injected with *cdon* MO at 256-cell stage (15.1%, n=92); c5,normal-loop in wild type controls (97.2%, n=72); c6-c8,wild-type embryos injected with *cdon* MO at 256-cell stage were examined for cardiac looping at 72hpf by WISH against *cmlc2* (n=110). **(D)** Percentages of normal looping, no looping, and reversed looping of the heart in all the embryos injected with or without *cdon* MO at 256-cell stage**. (E-F)** Expression of *Nodal/spaw* in controls (n=79) and embryos injected with *cdon* MO at the 256-cell stage (n=92). **(G-H)** Expression of *lefty1* in controls (n=86) and embryos injected with *cdon* MO at the 256-cell stage (n=91). **(I-J)** Expression of *lefty2* in controls (n=71) and embryos injected with *cdon* MO at the 256-cell stage (n=87). Notice: in “B” and “D’, all the transgenic embryos and wild type embryos were used together to calculate the percentage.

## 4. Discussion

Mouse Node/cilia (referred to as the ‘left-right organizer’ (LRO)) was first identified to play critical role during organ LR patterning (Kajikawa et al., 2022; Schneider et al., 1999; Sulik et al., 1994). Being similar to mouse, the transient structures were identified in other vertebrate embryos such as zebrafish, suggesting a conserved cilia-based mechanism regulates LR patterning in zebrafish (Essner et al., 2002). Indeed, functional studies confirmed the presence of motile cilia and asymmetric fluid flow in Kupffer’s vesicle in zebrafish (Essner et al., 2005) and genetic or embryological perturbation of these ciliated structures disrupted asymmetric Nodal pathway expression and organ laterality (Amack, 2014; Essner et al., 2005; Kuhns et al., 2019; Liu et al., 2019). In zebrafish, the transgenic line *Tg(sox17:EGFP)* has been developed to label the DFC/KV cells (Liu et al., 2019), and several developmental steps have been identified to build a functional KV/Cilia. DFCs appear at mid-epiboly stages, migrate, proliferate, and then undergo a mesenchymal-to-epithelial transition to form KV in early somite stage (Forrest et al., 2022). KV develops directional fluid flow and establishes LR signaling, and then breaks down around 18 hpf when KV cells undergo a epithelial to mesenchymal transition (Amack, 2021) and migrate away to incorporate into muscle and notochord (Ikeda et al., 2022). In the past decades, much of genes or environmental elements had been reported to involve in DFCs clustering migration (Ablooglu et al., 2010; Gao et al., 2011; Kajikawa et al., 2022; Lai et al., 2012; Liu et al., 2022), proliferation (Abdel-Razek et al., 2023; Gokey et al., 2015; Liu et al., 2019; Zhang et al., 2012) and the final KV formation and ciliogenesis, while the mechanisms underlying this process are from being completely elucidated.

Cdon is cell surface glycoproteins that belong to a subgroup of the immunoglobulin (Ig) super-family of cell adhesion molecules (Sanchez-Arrones et al., 2012). The role of Cdon in organ development and function had been reported in many literatures, including the role in neural differentiation, migration and survival (Jeong et al., 2014; Kim et al., 2023; Powell et al., 2015; Uluca et al., 2022; Wang et al., 2017), cardiac remodeling and fibrosis (Jeong et al., 2017) and myoblast fusion (Castiglioni et al., 2018). More recently, in zebrafish *cdon* was reported being involved in trunk neural crest cell migration and slow-twitch muscle development (Lencer et al., 2023) and limb growth (Echevarria-Andino et al., 2023). While in more early stage whether it plays a critical role in organ LR patterning is not reported.

Here, we identified *cdon* was expressed in DFCs during gastrulation movement (Fig. 1Aa2-a5) and in epithelial cells of KV at early somitogenesis stage (Fig. 1Aa6-a7). Further data showed that *cdon* loss of function led to DFCs clustering movement defect. As the results, KV formation/ciliogenesis and the subsequent organ LR patterning were disturbed. So, during embryonic development, beside the role in regulating trunk neural crest cell migration (Lencer et al., 2023) and defining the correct proximo-distal patterning of the eye development (Jeong et al., 2014), we additionally identified a more early role of *cdon* in regulating LR patterning. Comparing our data in *cdon* morphants and *cdon* mutants, we discovered that the organ LR patterning defect in *cdon* morphants is stronger than that in *cdon* mutants, the possible reasons are that: 1, in *cdon* mutant embryos, there still exists some degree of maternal *cdon* mRNA or matermal Cdon protein. 2, given the well-known genetic compensation response in zebrafish (Ma et al., 2019), the mutation of *cdon* possiblely upregulates some other genes to compensate the loss of *cdon* function. All these possibility need far more work to elucidate.

## Supplementary Materials

Figure s1. The expression of *boc* at different developmental stages

Figure s2. The external phenotype in embryos injected with *cdon* MO.

Figure s3. The external phenotype in embryos injected with *cdon* mRNA.

Figure s4. The external phenotype and the expression of *cdon* in controls and *cdon^-/-^*embryos.

Figure s5. The expression of *lefty1* and *lefty2* in controls and *cdon^-/-^*embryos.

Figure s6. Injection of *cdon* MO at 256-cell stage disturbed clustering DFCs migration, KV morphogenesis and ciliogenesis.

## Author Contributions

Conceptualization, supervision, and funding acquisition, S.H.; methodology, Z.D., Z.G. and S.H.; experiments, Z.D., W.C., C.L., B.L., S.H., K.Z., J.H. and S.H.; software, Z.D., Z.G. and S.H.; validation, Z.D., S.H.; formal analysis, Z.D., W.C., Z.G. and S.H.; data curation, Z.D.; writing-original draft preparation, Z.D., Z.G., Q.R and S.H.; writing—review and editing, S.H., Z.G., Z.D. and X.L. All authors have read and agreed to the published version of the manuscript.

## Informed Consent Statement

Not applicable.

## Data Availability Statement

All the data was in the manuscript and supplementary materials.

## Acknowledgments

We would like to thank Dr. Lingfei Luo for his advice on this work; we also would like to thank the members working in our fish facility for their help taking care of all the fish lines in this study.

